# Functional enrichment of alternative splicing events with NEASE reveals insights into tissue identity and diseases

**DOI:** 10.1101/2021.07.14.452376

**Authors:** Zakaria Louadi, Maria L. Elkjaer, Melissa Klug, Chit T. Lio, Amit Fenn, Zsolt Illes, Dario Bongiovanni, Jan Baumbach, Tim Kacprowski, Markus List, Olga Tsoy

## Abstract

Alternative splicing (AS) is an important aspect of gene regulation. Nevertheless, its role in molecular processes and pathobiology is far from understood. A roadblock is that tools for the functional analysis of AS-set events are lacking. To mitigate this, we developed NEASE, a tool integrating pathways with protein-protein and domain-domain interactions to functionally characterize AS events. We show in four application cases how NEASE can identify pathways contributing to tissue identity and cell type development, and how it highlights splicing-related biomarkers. With a unique view on AS, NEASE generates unique and meaningful biological insights complementary to classical pathways analysis.

## Background

Alternative splicing (AS) boosts transcript diversity in human cells [1] and thus contributes to tissue identity [2], cell development [3], and pathology in, e.g., cardiomyopathy [4], muscular dystrophy [5] or autoimmune diseases [6]. It is estimated that up to 30% of disease-associated genetic variants affect splicing [7]. RNA sequencing technologies (RNA-seq) allow the quantification of different types of AS events and detect splicing abnormalities in disorders. However, RNA-seq utility is currently limited by our incomplete understanding of the functional role of specific exons or the transcripts they contribute to.

A major challenge in AS analysis is the functional interpretation of a set of events, including isoform switching events and differentially spliced exons. The usual approach is to perform gene set enrichment or overrepresentation analysis [8–10]. This approach treats all genes affected by AS equally, neglecting that some AS events may not be functionally relevant at the protein-level [11] or result from noise in the splicing machinery [12]. Furthermore, functional differences between protein isoforms remain uncertain in many cases. A promising strategy to identify relevant AS events is to focus on those that lead to meaningful changes in the protein structure. Recent studies have shown that AS has the potential to rewire protein-protein interactions by affecting the inclusion of domain families [13] and linear motifs [14] or by activating nonsense-mediated decay [15].

This motivated the creation of databases and tools that predict the consequences of individual AS events or isoform switches. IsoformSwitchAnalyzeR [16], tappAS [17], DoChaP [18] and Spada [19] support transcript-level (as opposed to exon-level) analysis to identify isoform switches and their impact on the translation and the resulting isoforms features, such as domains, motifs, and non-coding sites. Exon Ontology [20] and DIGGER [21] support exon-level analysis to identify exon skipping events and their possible impact on the protein structure and function. Spada and DIGGER further consider the impact of AS on protein-protein interactions.

Most existing tools allow investigating AS-driven changes in an explorative fashion but tools for systematic analysis of functional effects of AS are lacking. Exon Ontology performs statistical tests to identify enriched features within a set of skipped exons. One example are domain families affected by AS across proteins more frequently than expected. However, none of the existing tools offer a systems biology view to specifically highlight functional consequences of AS events.

To tackle these limitations, we developed the first tool for functional enrichment of AS events. NEASE (Network-based Enrichment method for AS Events) first detects protein domains affected by AS and then uses an integrated protein-protein interaction (PPI) and domain-domain interaction (DDI) network [21] to identify protein interaction partners likely affected by AS. Next, it employs an edge-level hypergeometric test for gene set overrepresentation analysis. This approach is new in the way genes are selected for the enrichment test. Rather than considering only differentially spliced or expressed genes, which is currently the most common strategy, NEASE uses network information to select genes that are likely affected in the interactome. This is also superior to a simple network enrichment analysis, as we consider only those edges for which an AS contribution seems relevant and for which false positive results are less likely. We evaluated NEASE using multiple datasets from both healthy and disease cohorts. We show that the NEASE approach complements gene-level enrichment, and even outperforms it in scenarios where gene-level enrichment fails to find relevant pathways. Moreover, NEASE generates unique and meaningful biological insights on the exact impact of AS. Furthermore, since the statistical approach is network-based, NEASE can prioritize (differentially) spliced genes and finds new disease biomarkers candidates in case of aberrant splicing. The NEASE Python package, freely available at https://github.com/louadi/NEASE, provides multiple functions for a deeper analysis and visualization of affected protein domains, edges, and pathways (individually or as a set).

## Results

### Overview of NEASE

NEASE uses a hybrid approach that combines biological pathways with PPIs and DDIs to perform functional enrichment of AS. First, we use the structural annotation of known isoforms by mapping protein domains from the Pfam database [22] to the corresponding exons (Figure 1A). Second, we construct a structural joint graph as previously reported [21] by enriching the BioGrid PPI [23] with DDIs from DOMINE [24] and 3did [25] (see Methods). In the joint graph, protein domains are mapped to their mediated interactions. Thus, NEASE addresses the limited exon-level annotation and provides an exon-centric view of the interactome, where exons are represented by their encoded domains and edges represent the binding between the domains. In this way, the impact of AS can be seen as an edgetic change in the network. Analyzed AS events are viewed as a set of affected edges that represent gained or lost PPIs.

**Figure 1.**
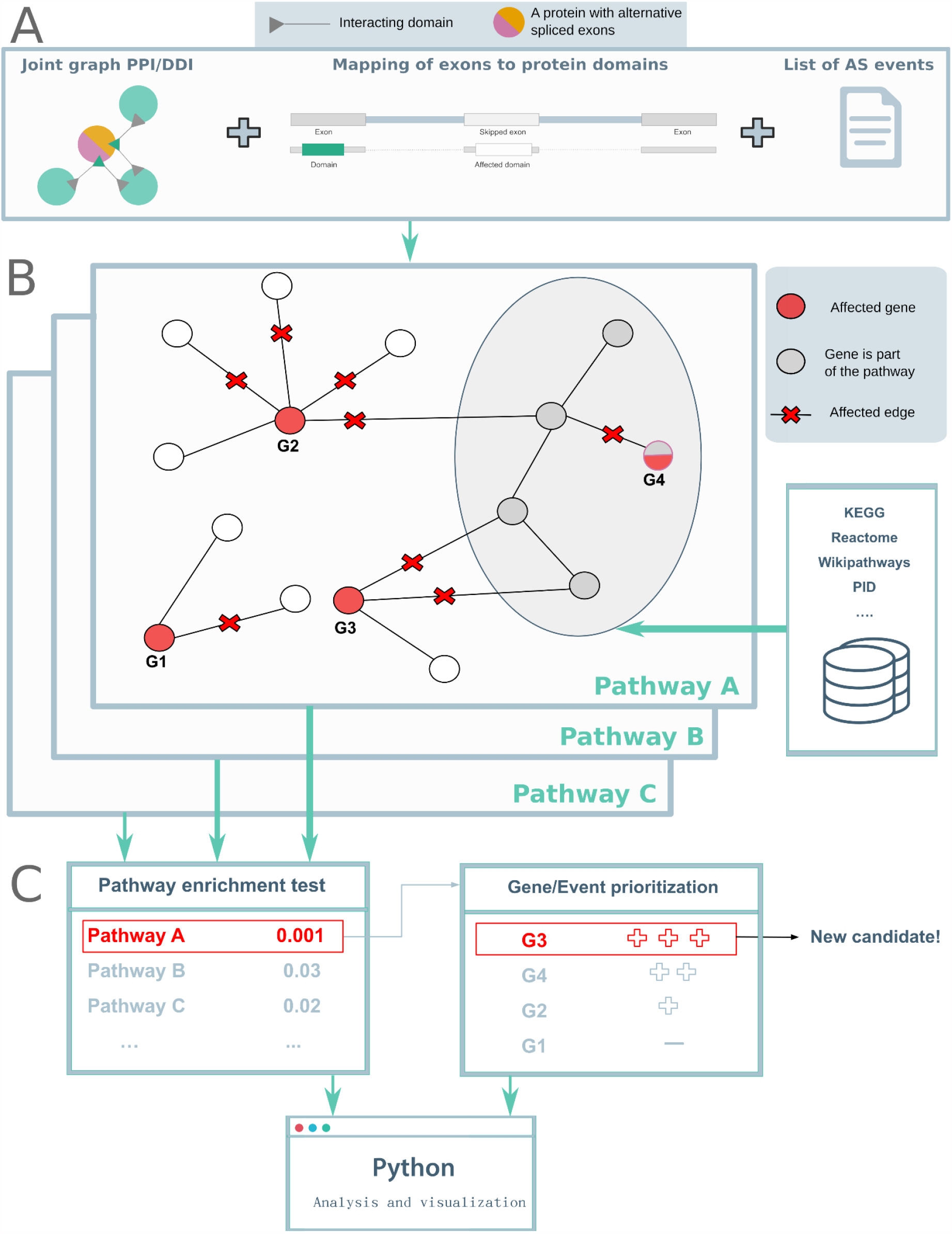
Overview of NEASE. (A) Annotated exons are mapped to Pfam domains. The joint graph of PPIs and DDIs is used to identify the interactions mediated by these domains. (B) For a list of exons/events, NEASE identifies interactions mediated by the spliced domains and pathways that are significantly affected by those interactions. (C) NEASE provides a corrected p-value, in addition to an enrichment score (NEASE Score) for every pathway (see Methods). The user can further focus on an individual pathway, where NEASE can prioritize genes and find new biomarkers. In this example, the gene G3 was not part of the enriched pathway A but it has the largest number of affected interactions with genes from the pathway.

We then perform statistical tests to find enriched pathways and most likely responsible genes (Figure 1B). Following, (differential) splicing analysis, a one-sided hypergeometric test is used to test for enrichment of a given pathway or gene set by considering all edges affected by AS in an experiment. A similar test is applied for each spliced gene to prioritize the most relevant events/genes that are affecting a pathway. We further introduce a weighted score (NEASE score) that penalizes hub nodes that are more likely to be connected to the pathway of interest by chance. Notably, this approach also considers genes that are not part of the existing pathway definition but show a significant number of interactions with the pathway, highlighting new putative biomarkers (see Methods for details).

The Python package provides an interactive analysis. Using a list of exons or events, users can run a general enrichment on 12 different pathway databases (collected from the ConsensusPathDB resource [26]), followed up by a specific analysis and visualization for a single affected pathway or module of interest (Figure 1C). To provide analysis for individual isoforms and events, we linked NEASE to our previously developed database DIGGER, which provides an isoform- and exon-centric view of the interactome [21].

### NEASE gives insights into the role of the muscle- and neural-specific exons

Recent studies suggest that the regulation of AS occurs in a tissue-specific manner and leads to remodeling of protein-protein interactions [27]. Understanding the functional impact of co-regulated exons is critical in understanding gene regulation. We applied NEASE to tissue-specific exons reported in VastDB, a resource that provides information on multiple types of AS events detected by RNA-seq from different tissue types and developmental stages [28]. We extracted 2,831 exon skipping events and Percent Spliced In values (PSI) from 12 different human tissue types (see Methods). We then performed hierarchical clustering on the z-score standardized PSI values (Figure 2A). The heatmap shows two distinct clusters, where neural-specific and muscle-specific (merged with heart-specific) exons are dominant.

**Figure 2.**
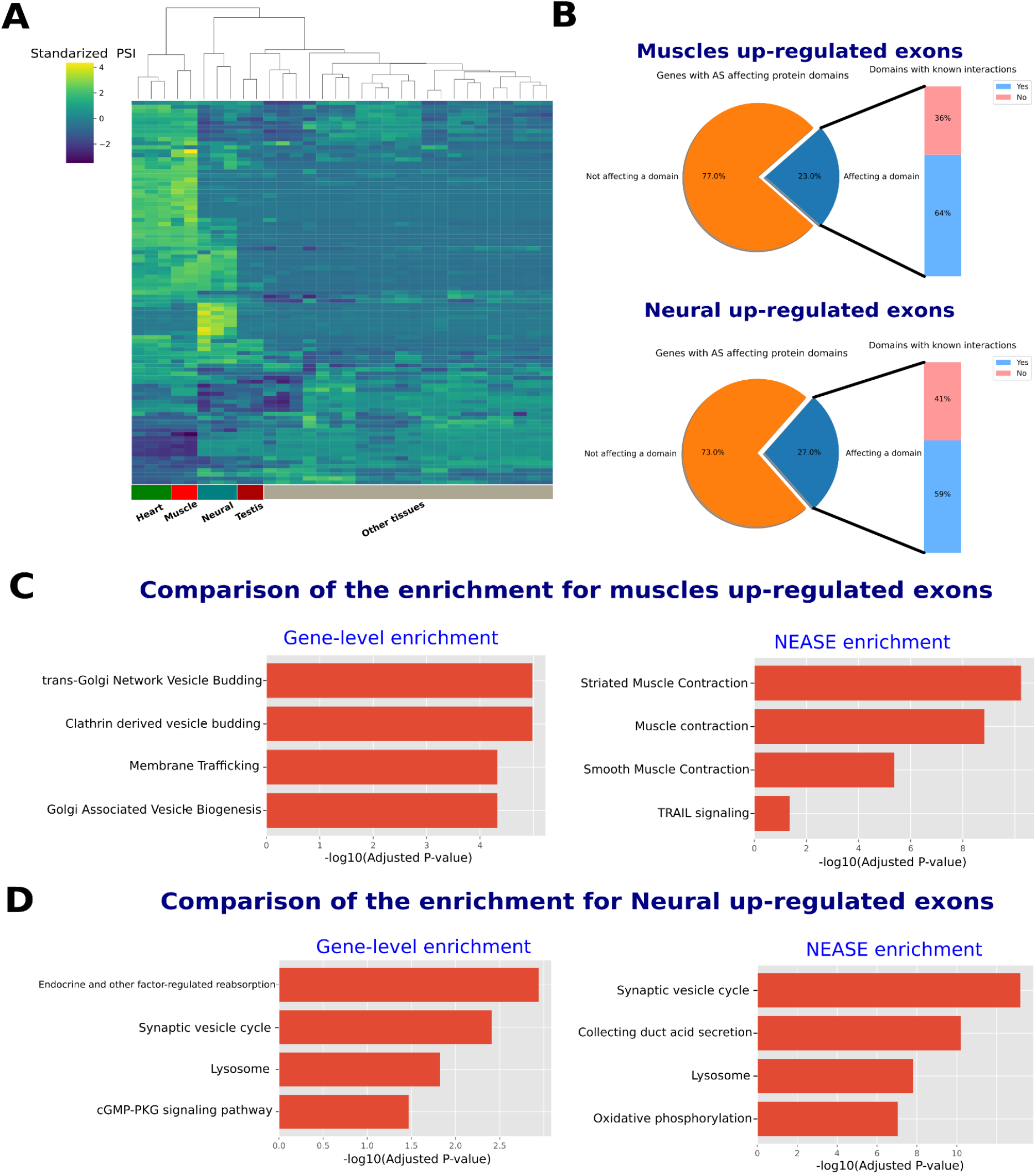
Analysis of tissue-specific exons. (A) Heatmap and hierarchical clustering of standardized PSI values obtained from VastDB. The heatmap only shows events with standard deviation of PSI values >=20. The heatmap shows that clusters of exons up-regulated in neural tissues and muscle/heart tissues are dominant. (B) NEASE analysis shows that 23% and 27% for both neural and muscle up-regulated exons, respectively, are encoding protein domains. Around 60% of the spliced domains are known to have interaction partners from the PPI and DDI joint graph. (C and D) Comparison between gene-level enrichment and NEASE enrichment for the two sets of exons.

Next, we extracted 66 skipped exons with a high PSI in the muscle tissues and 55 skipped exons with a high PSI in the neural tissues (z-score >= +2, see Methods). We checked how many of these events are overlapping with domains. As shown in Figure 2B, 23% of the upregulated exons in muscle tissues (13) and 27% of the upregulated exons in the neural tissues (17) overlapped with domains. NEASE also provides statistics of how many of these domains have known binding partners in the joint graph. In the two sets, around 60% of the affected domains have known interactions in our joint graph: 9 binding domains in the muscle tissues and 10 binding domains in the neural tissues (Additional files 2: Tables S4, S5 and Additional file 3: Tables S8, S9). For these groups of events, the exact protein complexes involved can be identified, and NEASE statistical analysis can be performed to determine affected pathways.

First, we ran a gene set overrepresentation analysis (one-sided hypergeometric test), which we refer to as gene-level enrichment, to detect enriched pathways (see Methods). Next, we applied NEASE to the same genes to detect pathways affected by AS. Unlike the gene-level enrichment, the results obtained from NEASE in both sets better explain the functional role of the regulated exons (Figure 2 C and D).

The upregulated exons in heart and muscle tissues were enriched in “Muscle Contraction” pathways (Figure 2C and Additional file 2: Table S7), while, in the gene-level enrichment, the pathways were related to very common subcellular functions such as the Golgi apparatus, which also is an organelle for collecting, modifying or destroying protein products (Figure 2 C and Additional file 2: Table S6). NEASE provides detailed information about the affected domains and their interaction partners (Additional file 1: Table S1). The domain Tropomyosin (Pfam id: PF00261), which is part of the gene TPM1, e.g., is involved in the regulation of muscle contraction via actin and myosin. GAS2 (Pfam id: PF02187) is a domain of DST, a dystonin encoding gene, which plays a role in maintaining the integrity of the cytoskeleton. AS affects its binding with the gene CALM1 that encodes a calcium-binding protein involved in various calcium-dependent pathways like muscle contraction [29].

The exons upregulated in neural tissues showed enrichment in the synaptic vesicle cycle pathway responsible for the communication between neurons (Figure 2D). Gene-level enrichment performed on par with NEASE, resulting in the same pathway but with a lower rank and significance (adjusted p-values: 6.51e-14 using nease and 0.0039 using gene-level, Additional file 3: Tables S10 and S11). Notably, NEASE also detected an enrichment in “oxidative phosphorylation”, which is the initiator for powering all major mechanisms mediating brain information and processing [30]. The neuron’s energy demands are remarkable both in their intensity and in their dynamic range and quick changes [31–34]. Therefore, AS could modify oxidative phosphorylation to serve the tissue-specific needs. Experimental studies have also found that several key enzymes in “oxidative phosphorylation” are spliced, e.g. pyruvate kinase (PKM) that shifts from the PKM2 to the PKM1 isoform [35,36]. NEASE also provides a detailed view on the affected mechanisms, such as an exon skipping event in the gene ATP6V0A1 overlapping with the V_ATPase_I domain (PFAM id: PF01496) and affecting the binding with seven other proteins from the complex vacuolar ATPase (V-ATPase) (p-value: 7.14e-17, Figure 3, Additional files 1: Table S2). V-ATPase is required for synaptic vesicle exocytosis [37] The a1-subunit of the V0 domain in ATP6V0A1 was recently shown to be highly expressed in neurons and to be essential for human brain development [38,39]. In another example, NEASE identified two co-regulated events of the genes CLTA and CLTB (Figure 3). CLTA and CLTB genes are involved in Clathrin-dependent endocytosis which forms clathrin-coated vesicles. Both genes play a major role in forming the protein complex of the coated vesicle. Both events affect the same domain Clathrin light chain (Pfam id: PF01086). The Clathrin light chain domain binds to CLTC and CLTCL1 which are the Clathrin heavy chain genes (p-value: 7.356981e-05).

**Figure 3:**
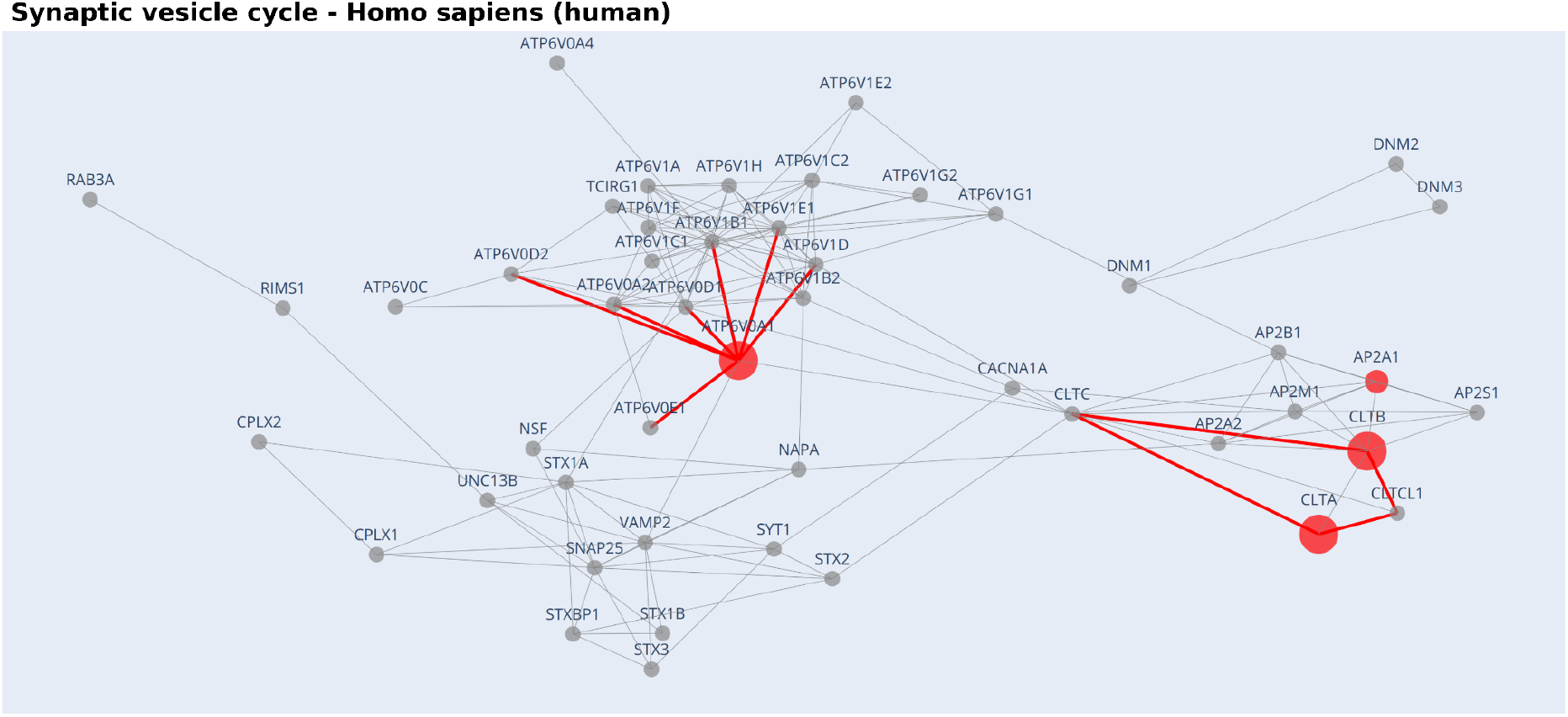
NEASE visually highlights the impact of the AS regulation at the interactome level. The gray nodes represent proteins from the pathway and the red nodes represent genes with AS events. Red edges represent the affected interactions for the nodes with known DDIs. The visualization of the pathway “Synaptic vesicle cycle” from the KEGG database for the exons up-regulated in the neural tissues shows that the splicing in the genes CLTA and CLTB is co-regulated and affects the interactions of the same complex. Similarly, NEASE highlights the importance of the domain ATP6V0A1 which is up-regulated in neural tissues and binds seven proteins from the “Synaptic vesicle cycle” pathway.

These results suggest that the formation of this complex is co-regulated by AS. A similar finding about the role of the Clathrin light chain in neurons was also described in [40]. NEASE highlights these co-regulated events at the network level (Figure 3). The analysis generated from VastDB using NEASE agrees with the latest studies at transcriptomics and proteomics levels that emphasize the crucial role of AS in the function and development of brain and heart tissues [41–43].

### NEASE reveals splicing-related differences of reticulated and mature platelets

AS does not only drive tissue-specific regulation but also plays a major role in cell differentiation and maturation. To illustrate an example of the utility of NEASE in such studies, we used the RNA-seq data set from [44] which compares the transcriptome profiles of reticulated platelets and mature platelets from healthy donors. Reticulated platelets are younger [45], larger in size, and contain more RNA [46]. Moreover, they have a prothrombotic potential and are known to be more abundant in patients with diabetes, acute or chronic coronary syndrome, and in smokers [46–48]. Additionally, elevated levels of reticulated platelets in peripheral blood are predictors of insufficient response to antiplatelet therapies (e.g. Aspirin and P2Y12 inhibitors) and are promising novel biomarkers for the prediction of adverse cardiovascular events in different pathological settings [47,49]. A strong enrichment of pro-thrombotic signaling in reticulated platelets was observed in healthy donors [44]. Comparative transcriptomic analysis revealed a differential expression of several pathways in addition to an enrichment of prothrombotic pathways and transcripts of transmembrane proteins as the collagen receptor GPVI, the thromboxane receptor A2 and the thrombin receptors PAR1 and PAR4. Gene set enrichment analysis indicated an upregulation of entire prothrombotic activation pathways as the thrombin PAR1 and integrin GPIIb/IIIa signaling pathway in reticulated platelets.

Since AS has been described to occur in platelets [50], we wanted to investigate the splicing patterns between the previously defined reticulated and mature platelet subgroups. Using MAJIQ [51] (see Methods), we found 169 differentially spliced genes. From 25 affected protein domains, 17 have known interactions (68% of affected domains, Figure 4 A, Additional file 4: Tables S12 and S13).

**Figure 4.**
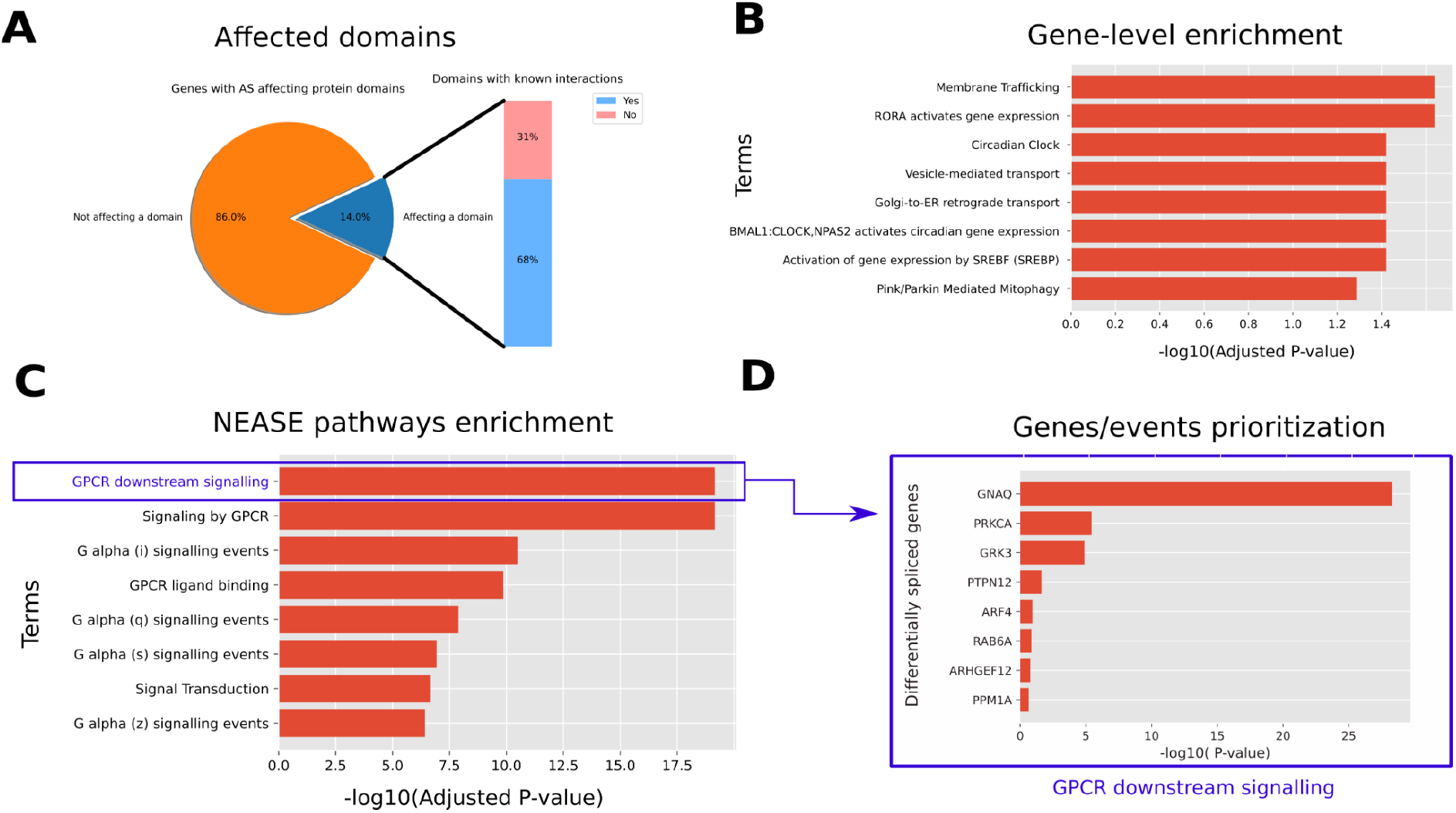
(A) 14 % of differentially spliced exons, between reticulated and mature platelets, are known to encode protein domains. (B) Gene level enrichment of all differentially spliced exons in the Reactome database fails to capture most relevant pathways. (C) In contrast, NEASE shows an enrichment of the GPCR downstream signaling and other related pathways that are well known to be important in platelet activation. (D) A further look at the genes driving the enrichment of the GPCR pathway shows the most relevant genes affected by AS.

We observed that the enrichment at the gene-level using the Reactome [52] database ranks general cellular pathways higher, including “Membrane Trafficking” and “Vesicle-mediated transport”, and “Golgi-to-ER retrograde transport”. An exception is the “Circadian Clock” pathway, which is hypothesized to be related to platelet activation [53] (Figure 4 B). The pathway “Platelet activation, signaling and aggregation” was less significant in gene-level enrichment (adjusted p-value: 0.061, Additional file 4: Table S14) compared to NEASE enrichment (adjusted p-value: 0.004, Additional file 4: Table S15). Using NEASE, we obtained more meaningful results and unique pathways. As shown in Figure 4C, the most significant pathways in reticulated platelets are G Protein-Coupled Receptor-related. G proteins are essential in the second phase of platelet-dependent thrombus formation [54]. Furthermore, GPCR isoforms are known to have distinct signaling properties [55]. Other relevant pathways associated with platelet activation are “Hemostasis”, “Thromboxane signaling through tp receptor”, and “Platelet homeostasis”. The full tables for enrichment at the gene level and using NEASE are available in the Additional file 4: Tables S14 and S15. The upregulation of these pathways in reticulated platelets emphasizes their previously described prothrombotic phenotype and their involvement in several downstream signaling processes.

We also looked at the individual AS events driving this enrichment. For each affected domain, NEASE tests if it significantly interacts with the GPCR downstream signaling pathway (Additional file 4: Table S16, see Methods). Figure 4C illustrates affected genes and their p-value ranking. The top gene is GNAQ (G-protein subunit alpha q), which is known to be involved in signal transduction in platelets leading to platelet activation [56]. The regulation of the G-protein alpha subunit can be an indication that compared to mature platelets, reticulated platelets are more involved in various signal transduction pathways related to, e.g., pro-thrombotic processes [46]. PRKCA, which also showed different splicing patterns between the two platelet subgroups, plays a major role in the platelet formation process by modulating platelet function [57], megakaryocyte function, and development [58] and negatively regulates pro-platelet formation [59]. Moreover, the regulation of PRKCA binding in reticulated platelets might refer to the young nature of reticulated platelets, which have undergone the pro-platelet formation process more recently than mature platelets [45,60].

### NEASE characterizes complex disorders such as Multiple Sclerosis

Multiple sclerosis (MS) is a chronic inflammatory demyelinating disease of the central nervous system. Early in the disease course, MS is characterized by focal lesions in the brain induced by influx of systemic inflammatory cells. These active lesions infiltrated by immune cells and activated microglia are characterized by inflammatory demyelination and axonal loss [61]. The surrounding white matter tissue is termed normal-appearing white matter due to diffuse pathology without focal lesion activity and dense immune activity [62]. The etiology of MS remains unknown. Recently, a systematic literature review found 27 genes that were alternatively spliced in MS patients [63].

We used RNA-Seq of macrodissected areas from postmortem white matter tissue of patients with progressive MS [64]. We compared normal-appearing white matter and active lesions regions from postmortem white matter brains of MS patients. We found 109 differentially spliced genes and 19 affected domains with known interactions. In total, NEASE identified 150 affected interactions (Additional file 5: Tables S17 and S18).

Gene-level enrichment ranks high pathways likely irrelevant that are involved in muscle contraction, cardiac conduction, and membrane Trafficking, with the exception of Ca2+ ion flow across membranes.(Additional file 5: Table S19). Ca2+ is an essential signal molecule for all cell activity. Although deregulation of calcium signaling is related to the pathogenesis of multiple diseases [65], including neurological disorders [66], It is not specific to neuronal tissues. In line with the neurodegenerative and immune-mediated features of MS, NEASE found unique enriched pathways related to brain network signaling and neuronal pathways “Neurotransmitter receptors and postsynaptic signal transmission”, “Transmission across Chemical Synapses”, “Activation of NMDA receptor and postsynaptic events”, “MAPK family signaling cascades”, “Neuronal System”), as well as pathways related to immune responses (“interleukin-17 signaling, “Toll Like Receptor 10 (TLF10) Cascade”) (Table 1 and Additional file 5: Table S20). Two other pathways were related to the uptake of anthrax or bacterial toxins. This could be a result of clean-up from toxic inflammatory processes or increased presence of invaders due to the leaky brain-blood-barrier in MS [67–69]. Additionally, it also supports the theory of infections as the trigger of lesion damage in MS [70].

**Table 1.** NEASE enrichment obtained from AS comparison between normal-appearing white matter (NAWM) and acute lesions (AL), from multiple sclerosis patients. The highly enriched pathways belong to Neurotransmitter receptors, MAPK, and bacterial infection. Most of these pathways are hallmarks of MS. The NEASE score is obtained after combining the p-value with the number of significant genes. The latter is obtained after individual tests for each gene in the column “Spliced genes” (see Methods).

| Pathway name | Spliced genes (number of interactions affecting the pathway) | p-value | adj p-value | NEASE score |
| --- | --- | --- | --- | --- |
| Neurotransmitter receptors and postsynaptic signal transmission | GRIA1 (7), ATP2B1 (1), BRAF (4), MAP2K4 (1), GRIN1 (4) | 2.128010e-08 | 0.000009 | 15.34 |
| Uptake and function of anthrax toxins | ATP2B1 (1), BRAF (5), MAP2K4 (3) | 2.468257e-09 | 0.000004 | 14.90 |
| Transmission across Chemical Synapses | GRIA1 (7), ATP2B1 (1), BRAF (4), MAP2K4 (1), GRIN1 (4) | 2.207774e-07 | 0.000037 | 13.31 |
| Uptake and actions of bacterial toxins | ATP2B1 (1), BRAF (5), MAP2K4 (3) | 2.560786e-08 | 0.000009 | 13.14 |
| Activation of NMDA receptor and postsynaptic events | GRIA1 (2), ATP2B1 (1), BRAF (4), MAP2K4 (1), GRIN1 (3) | 2.447069e-06 | 0.000194 | 11.22 |
| MAPK family signaling cascades | MYH10 (2), ATP2B1 (1), BRAF (17), MAP2K4 (5), GRIN1 (3) | 1.140595e-06 | 0.000148 | 10.29 |
| Neuronal System | GRIA1 (7), ATP2B1 (1), BRAF (4), MAP2K4 (1), GRIN1 (4) | 2.657946e-06 | 0.000194 | 9.65 |
| FCERI mediated MAPK activation | MYH10 (1), BRAF (7), MAP2K4 (8) | 5.455227e-07 | 0.000082 | 8.85 |
| RAF/MAP kinase cascade | MYH10 (1), ATP2B1 (1), BRAF (16), MAP2K4 (4), GRIN1 (3) | 7.076613e-07 | 0.000099 | 8.69 |
| MAPK1/MAPK3 signaling | MYH10 (1), ATP2B1 (1), BRAF (16), MAP2K4 (4), GRIN1 (3) | 1.736848e-06 | 0.000194 | 8.14 |
| Interleukin-17 signaling | BRAF (6), MAP2K4 (12) | 2.203770e-06 | 0.000194 | 7.99 |
| MAP kinase activation | BRAF (6), MAP2K4 (12) | 2.203770e-06 | 0.000194 | 7.99 |
| Toll Like Receptor 10 (TLR10) Cascade | BRAF (6), MAP2K4 (13) | 2.618844e-06 | 0.000194 | 7.89 |

As shown in Table 1, the pathway “Uptake and function of anthrax toxins” has the best overall adjusted p-value, calculated only based on the total number of edges affecting the pathway. When we also included the number of significant genes and calculated NEASE scores (see Methods), NEASE ranks the pathway “Neurotransmitter receptors and postsynaptic signal transmission” first, and moves pathways such as “Transmission across Chemical Synapses’ and “Neuronal System” higher in the rank. These observations illustrate the usefulness of the NEASE score as a complement to the global edge-based enrichment.

Two of the most significant genes in the “Neurotransmitter receptors’’ pathway were GRIN1 and GRIA1 (Additional file 5: Table S21). GRIN1 encodes GluN1, which is one of the two obligatory subunits for the NMDAR1 receptor, where GRIA1 encodes the AMPAR1 subunit. Their ligand is glutamate, and they are both ionotropic receptors and have been associated with MS disease severity [71–73]. Interestingly, AS of MAP2K4 appeared in both brain-related and immune-related pathways, significantly enriched in active lesions *vs* normal-appearing white matter (Table 1). MAP2K4 is a mitogen-activated protein kinase (MAPK) orchestrating multiple biological functions [74,75]. AS of MAP2K4 has been found in rheumatoid arthritis [76], as well as in pathways of patients with other autoimmune diseases [77]. MS also precedes autoimmune attack, and therefore AS of MAP2K4 in active lesions detected with NEASE may represent dysregulated immune responses originating from the infiltrating immune cells or inflammatory-activated brain cells. This is supported by previous studies that found (i) overactivity of MAPK pathways in microglia (the resident immune cell of the brain) during neurodegeneration [78,79], and *(ii)* increased phosphorylation of MAPK kinases in the systemic immune cells of MS patients [80,81]. A recent study also characterized activated MS-specific pathways in immune cells from blood using phosphoproteomics. Here, MAP2K4 and its interaction partners (e.g. TAK1) were present in MS-specific signaling activity [82]. Future functional studies on the AS of MAP2K4 may help explain if AS could be the reason for increased phosphorylation and overactivity detected in MS. AS of MAP2K4 could result in switching protein conformation, increasing susceptibility to phosphorylation or changing the downstream protein cascade.

With NEASE, we were able to specifically detect AS of genes and related pathways already known to be dysregulated within MS from excitotoxicity to inflammation. The detected AS genes in active lesions *vs* normal-appearing white matter demonstrate how major components in signaling activities may be fine-tuned/changed from regulation of a homeostatic state to an inflammatory state. Combining NEASE with functional experiments to understand the biological impact of AS could fuel new therapeutic opportunities for complex neurological diseases as MS. Novel developments in genome-editing tools and gene-specific strategies have made it possible to use antisense oligonucleotides or small modulators for splice modification. This is already used in the rare neuromuscular disease, spinal muscular atrophy, where an antisense oligonucleotide binds to a site near splicing to ensure the inclusion of an exon during the splicing event [78].

### NEASE finds new biomarker candidates for Dilated Cardiomyopathy

AS might play a role in driving Dilated Cardiomyopathy (DCM) [83]. DCM is a common heart muscle disease that is often diagnosed with structural abnormalities resulting in impaired contraction. Previous studies have shown a large number of differentially used exons in DCM patients [4,10]. In this analysis, we used a list of 1,212 differentially used exons between DCM patients and controls as reported by Heinig *et al*. [10]. 28% of these exons overlapped with domains (Additional file 6: Tables S22 and S23). In this exon set, both the gene level enrichment and NEASE show very similar results (Additional file 6: Tables S24 and S25). In both methods, we found that the list of exons was enriched in the Dilated cardiomyopathy (DCM) pathway from KEGG, as well as, “Adrenergic signaling in cardiomyocytes”, and “Regulation of actin cytoskeleton”.

In contrast to gene-level enrichment analysis, NEASE is able to score the contribution of alternatively spliced genes that are interacting with but are not part of the DCM pathway, allowing us to highlight putative biomarkers (Table 2, Additional file 6: Table S26, Additional files 1: Figure S1). The Myosin head domain from the gene MYO19 interacts with 6 other genes associated with DCM: (1) MYL2, which triggers contraction after Ca+ activation [84]; (2-5) TPM1/TPM2/TPM3/TPM4, which encode the tpm protein - the main regulator of muscle contraction [85]; and (6) ACTG, which encodes actin. Interestingly, MYO19 has not been investigated for its role in DCM, while its interacting genes are associated with DCM [86–89]. Additionally, the gene OBSCN has one affected interaction with the TTN gene [90]. The TTN gene itself is also differentially spliced associated with DCM [90]. OBSCN was recently reported as a new DCM candidate [91,92]. Another interesting example is CACNA1C (Calcium Voltage-Gated Channel Subunit Alpha1 C), an already known DCM candidate [93]. The differentially spliced exon overlaps with the domain Ion_trans (Pfam id: PF00520) that is essential for myocyte contraction [94]. The affected interaction identified is with the Ryanodine receptor 2 (RYR2). In striated muscles, excitation-contraction coupling is mediated by this complex [95]. Both CACNA1C and RYR2 are part of the KEGG DCM pathway [96]. Alterations in Ryanodine Receptors were repeatedly reported to be related to heart failure [97–99].

**Table 2:**
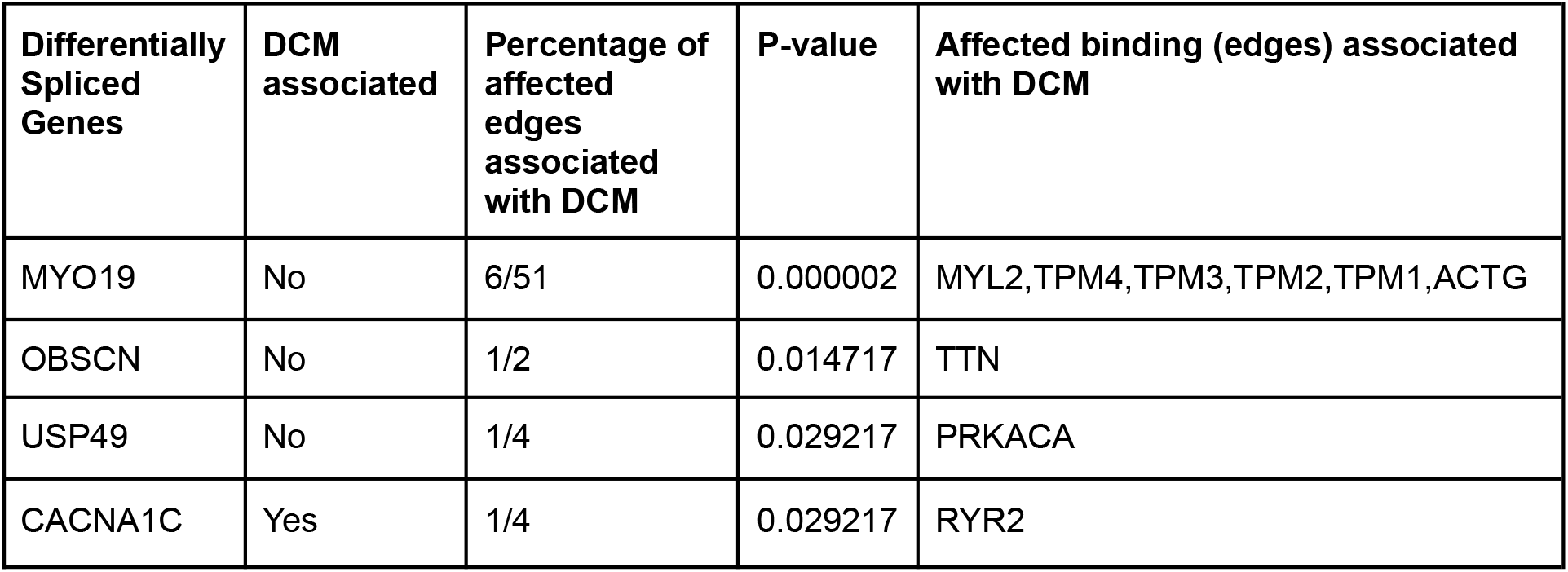
Enrichment of the pathway “Dilated cardiomyopathy (DCM)” from KEGG for the exons differentially used in DCM patients. The table shows the most significant genes (p_value<0.05) (See Methods).

## Discussion

In spite of its importance for biomarker and therapeutic target discovery, differential AS is still not a routine part of transcriptome analysis. A key reason for this could be the lack of suitable methods and software tools for AS-specific functional analysis. Our method NEASE closes this gap and provides a unique view on the impact of AS complementary to functional insights gained from traditional gene-level enrichment analysis. We applied NEASE to four diverse data sets and show that it’s results generate novel disease-relevant insights and provide valuable context to prior findings on altered RNA- and protein-expression levels consistent with recent literature.

In many cases, NEASE improves over gene-level enrichment analysis focusing on differentially spliced genes. One potential reason for this could be that not all AS events are necessarily functional [11,12]. NEASE mitigates this by focusing on AS events that affect protein domains. However, it is important to keep in mind that this is not the only way to define functional AS events. AS also affects interacting disordered regions [14] or facilitates nonsense-mediated decay [100].

AS events could also lead to completely different functions or interactions [101], e.g., two isoforms can have different interaction partners depending on the inclusion or loss of a single domain [13]. Such changes in the interactome can not be captured with gene-level enrichment which has a strict focus on nodes rather than edges. With NEASE, we could show that integrating structural information at the exon level and PPI networks helps to identify the functional impact of differentially spliced and co-regulated exons. In practice, we consider both approaches as complementary and recommend running gene-level and edge-level enrichments together (both supported by the NEASE package).

NEASE relies on structurally annotated interactions and existing pathway annotations from databases such as KEGG [96] and Reactome [52]. Leveraging reliable structural information and established pathways likely removes many false positive PPI from considerations. While DDI are generally of high quality, it should be noted that not all AS events are necessarily of functional consequence since other processes such as nonsense-mediated decay need to be considered as well. With our current approach, a large fraction of the PPI network remains unexplored, suggesting that adapting *de novo* network enrichment methods such as KeyPathwayMiner [102] towards AS could be a promising research direction to uncover previously unknown disease mechanisms. NEASE currently considers the immediate neighborhood of a pathway in the PPI network. When carefully considering the expected increase in false positives, one could also increase the size of the pathway neighborhood using, e.g., a fixed radius for shortest paths. While these are attractive approaches, the biases of the PPI towards hubs, as well as the high number of false (or missing) edges of PPI, in its current form, make such approaches hard to control and statistically challenging. Even though NEASE is relatively conservative, we demonstrated that it is simple, robust, and generates meaningful and interpretable results. Thus, it provides an unprecedented opportunity to understand the functional impact of tissue-, developmental- and disease-specific AS in a system biology manner.

While a plethora of gene set enrichment methods have been proposed in recent years, AS is typically not addressed specifically. Thus, NEASE closes an important gap in functional enrichment analysis of transcriptomics data. The analyses described here, confirm the widespread impact of AS in multiple biological processes and disorders. In the future, we plan to extend NEASE with further model organisms and to add structural annotations covering more types of AS events. Finally, we plan to integrate NEASE with the DIGGER web tool [21] for a seamless downstream analysis of AS in the web browser with the vision of establishing functional AS event analysis as a routine step in transcriptomic analysis.

## Methods

### NEASE Data sources

We construct a human structurally annotated PPI as described previously [21]. Briefly, we integrate DDI and PPI information into a joint network where DDIs were obtained from 3did (v2019_01 [25]) and DOMINE (v2.0 [24] including high- and mid-confidence interactions) and PPIs were obtained from BioGRid 3.5 [23]. In summary, out of 410,961 interactions from the human interactome 52,467 have at least one domain interaction. The mapping of exons to domains was performed using DIGGER’s mapping table that in turn uses BioMart and Pfam annotations [22,103]. We obtain the biological pathways with their gene list from KEGG [52] and Reactome [96] integrated into the ConsensusPathDB database [26].

### Statistical tests and pathway scores

Gene-level enrichment is performed using a hypergeometric test from the package GSEAPY (a Python wrapper for Enrichr [104]) by considering all genes with (differential) AS events.

For NEASE enrichment, we filtered the PPI graph G=(V,E), where V is the set of genes and E is the set of edges, to a subgraph G’=(V’, E’) containing only structurally annotated interactions E’ and their nodes V’.

For a submitted query list of exons, NEASE first identifies affected domains that overlap with the exons and their interactions. Let N be two times the number of edges in G’ (the degree of the network) and n be the number of affected edges from the query. These edges are then considered using a test modified from [105]: For every pathway P with degree K, let k be the number of affected edges that are connected to P. We model X whose outcome is k as a random variable following a hypergeometric distribution:

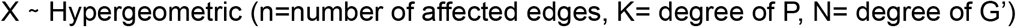

where k is considered as the number of observed successes out of n draws, from a population of size N containing K success. Subsequently, NEASE tests if the number k is significant using a one-sided hypergeometric test (over-representation). After testing for multiple pathways, the obtained p-values for the edge-level enrichment are corrected, using the Benjamini-Hochberg method [106].

For a pathway of interest, a similar test can be applied to determine if a splicing event significantly affects interactions of a specific gene with this pathway. Here, n is the number of all affected interactions (edges) of a spliced gene and k is the number of affected interactions (edges) across genes that are linked to the pathway of interest. As a result, for every pathway, NEASE provides an overall p-value, as well as the most significant genes. Since the p-value only depends on the overall number of affected edges but not on the number of genes, the p-value can be heavily influenced by hub genes. To reduce this influence, an optional score (NEASE Score) can be computed by NEASE to scale the natural logarithm of the p-value with the total number of significant genes using a cutoff from the user (for instance p-value<0.05):

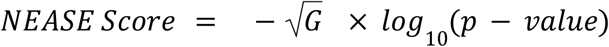

where G is the total number of significantly connected genes obtained after testing individual spliced genes. Thus, the NEASE Score prioritizes pathways that are affected by a larger number of spliced genes rather than pathways that have a larger number of affected interactions (edges). The user can choose to rank enrichment based on the adjusted p-value or by the NEASE score.

### VastDB events processing

PSI values of the exon skipping events from VastDB were quantified by the developers using vast tools [28,107]. In our analysis, we extracted the PSI values for 32 experiments belonging to 12 main tissues: muscle/heart, neural (whole brain, cortex and peripheral retina), placental, epithelial, digestive (colon and stomach), liver, kidney, adipose, testis, immune-hematopoietic and ovary. We then filtered out the events with low read coverage (VLOW) and performed hierarchical clustering of standardized values (z-scores). For every exon, we calculated the mean of PSI values from the samples of the same tissues. To extract muscle/heart and neural-specific exons, we applied two filters, namely that the z-score of the exon PSI value in the relevant tissue is higher than 2 and that the mean PSI value of the exon across all tissues is between 0.15 and 0.90. The latter ensures that we consider only exons that are part of AS events.

### RNA-Seq analysis

Raw RNA-Seq reads for two types of platelets and multiple sclerosis patients were downloaded from the GEO repository (access numbers: GSE126448 and GSE138614). The number of samples and sequencing depth are reported in Additional file 1: Table S3. RNA-Seq reads were aligned to the reference human genome (hg38) using STAR 2.7 [108] in a 2-pass mode and filtered for uniquely mapped reads. Differential AS analysis was performed by MAJIQ [51] with default parameters, and with a threshold of P(dPSI > 20%) > 0.95.

### NEASE: The Python package

NEASE’s Python package relies on NumPy [109], pandas [110], NetworkX [111], SciPy [112], and Statsmodels [113]. The gene-level enrichment is also supported in the NEASE package using the Python implementation of Enrichr [104]. To speed up the edge hypergeometric test, the total degree of every pathway in the structural PPI, as well as the overall degree of the network were pre-computed. For visualization, we use the complete PPI (not the structural PPI) and extract connected subnetworks from each pathway as well as spliced genes and their interactions with the extracted modules. The position of nodes is computed using the Fruchterman-Reingold force-directed algorithm implemented in NetworkX [114]. The interactive visualization for individual genes and events is implemented with information from the DIGGER database and the Plotly package.

The standard input of the package is a DataFrame object with the exon coordinates and Ensembl IDs of the genes. The package also supports the output of AS differential detection tools that are event-based such as MAJIQ [51] where NEASE only considers annotated exons. NEASE is released as open-source under the GPLv3 license and available at (https://github.com/louadi/NEASE). The code used to produce the results in this manuscript was deposited at Zenodo (https://doi.org/10.5281/zenodo.4985321).

## Supporting information

Additional file 2: upregulated exons in muscles

Additional file 3: upregulated exons in neural

Additional file 4: Differential splicing between reticulated platelets and mature platelets

Additional file 5: Differential splicing between normal-appearing white matter and acute lesion from multiple sclerosis patients.

additional file 6: Differential splicing between Dilated Cardiomyopathy patients and controls

## Declarations

### Availability of data and materials

RNA sequencing data for reticulated platelets was provided by the authors [44] and it is freely available at GEO (access number: GSE126448). Multiple sclerosis Raw sequence was provided by the authors [64] and freely available at GEO (access number: GSE138614). Dilated Cardiomyopathy raw data is available in the European Genome-phenome Archive (Dataset ID: EGAS00001002454), in our analysis we used pre-processed data from the manuscript [10]. Vastdb dataset for humans (hg19) was downloaded from https://vastdb.crg.eu/wiki/Downloads. The generated joint graph and the exon mapping database are available on the DIGGER database website https://exbio.wzw.tum.de/digger/download. All processed datasets, as well as step-by-step tutorials for using NEASE to reproduce the results presented in this paper, are available at https://github.com/louadi/NEASE-tutorials and deposited to Zenodo (https://doi.org/10.5281/zenodo.4985321).

### Authors’ contributions

ZL, TK, OT, JB and ML conceived the project. ZL performed the initial experiments, developed the method, and implemented the software. OT and ML supervised the project, provided critical feedback, and helped shape the research. OT prepared the RNA-Seq datasets for the differential splicing analysis. AF, CTL interpreted and discussed the biological insights of the results. MK and DB interpreted and discussed the platelet analysis. MLE and ZI interpreted and discussed the Multiple Sclerosis analysis. ZL wrote the initial draft of the manuscript. All authors contributed to writing the final manuscript and approved the final version.

## Acknowledgements

Not applicable.

