## Additional file 1: Supplemental figures and tables. for "Functional enrichment of alternative splicing events with NEASE reveals insights into tissue identity and diseases"

**Supplementary Table S1:** Enrichment of the pathway “Muscle contraction” from Reactome for the exons up-regulated in the muscles generated by the NEASE package.

| Spliced Genes | Affected domain | Gene is known to be in the pathway | Percentage of affected edges associated to the pathway | P-value | Affected binding (edges) |
|---|---|---|---|---|---|
| TPM1 | PF00261 | Yes | 10/18 | 8.30E-16 | TPM2, MYH8, TNNT2, MYH6, ACTA2, TPM4, TPM1, TNNT1 TPM3, TNNI1 |
| DST | PF02187 | No | 1/1 | 1.08E-02 | CALM1 |
| LIMS1 | PF00412 | No | 1/8 | 8.31E-02 | PXN |

**Supplementary Table S2:** Enrichment of the pathway “Synaptic vesicle cycle” from KEGG for the exons up-regulated in the neural tissues generated by the NEASE package.

| Spliced Genes | Affected domain | Gene is known to be in the pathway | Percentage of affected edges associated to the pathway | P value | Affected binding (edges) |
|---|---|---|---|---|---|
| ATP6V0A1 | PF01496 | Yes | 7/7 | 8.304262e-16 | ATP6V1B1, ATP6V0A2, ATP6V0D2, ATP6V1E1, ATP6V0E1, ATP6V0D1, ATP6V1 |
| CLTA | PF01086 | Yes | 2/3 | 7.356981e-05 | CLTC, CLTCL1 |
| CLTB | PF01086 | Yes | 2/6 | 3.642251e-04 | CLTC, CLTCL1 |

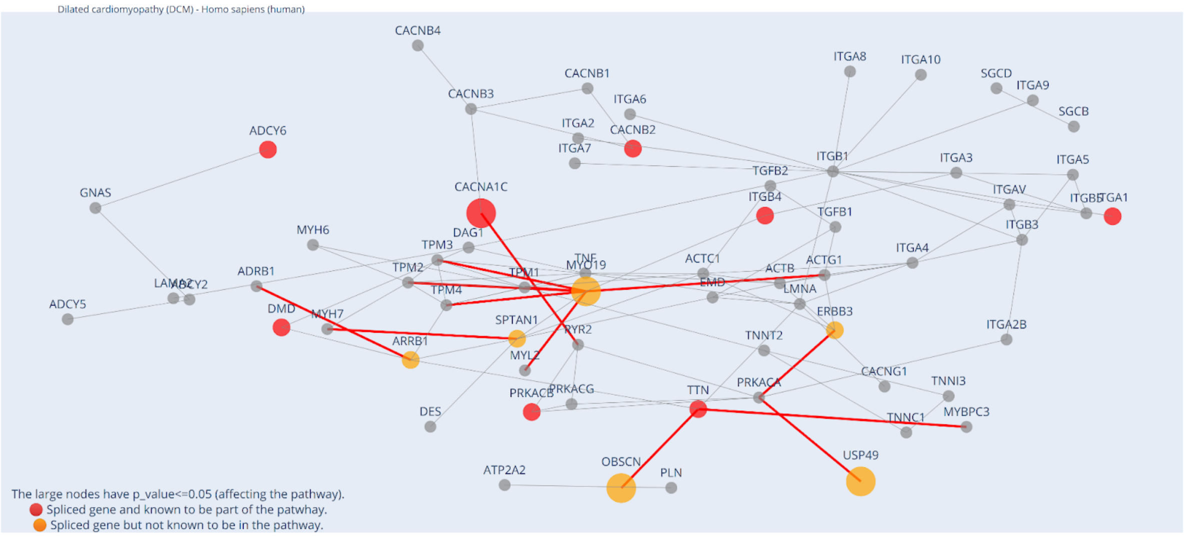


**Supplementary Figure S1**: NEASE visualization highlights the interactions of differentially spliced genes with the DCM pathway. The gray nodes represent proteins and the red nodes represent differentially spliced genes (both from the pathway). Orange nodes represent differentially spliced genes that are not in the DCM pathway but have affected interactions with the other genes from it. Red edges represent the affected interactions.

**Supplementary Table S3**. RNA-Seq samples used in the study

| Reticulated and mature platelets [1] | | |
|---|---|---|
| Description | Number of samples | Sequencing depth |
| Mature platelets | 4 | 15 million single reads |
| Reticulated platelets | 4 | 20 million single reads |
| Multiple sclerosis [2] | | |
| Normal-appearing white matter (NAWM) | 15 | 13 - 84 million paired reads |
| Acute lesion (AL) | 20 | 29 - 301 million paired reads |
| Dilated Cardiomyopathy (the used data is from [3]) | | |
| Heart samples of patients (DCM) | 97 | 90 - 246 million paired reads |
| Heart samples of healthy donors | 108 | 106 - 231 million paired reads |

1. Bongiovanni D, Santamaria G, Klug M, Santovito D, Felicetta A, Hristov M, et al. Transcriptome Analysis of Reticulated Platelets Reveals a Prothrombotic Profile. Thromb Haemost. 2019;119:1795–806.

2. Elkjaer ML, Frisch T, Reynolds R, Kacprowski T, Burton M, Kruse TA, et al. Molecular signature of different lesion types in the brain white matter of patients with progressive multiple sclerosis. Acta Neuropathol Commun. 2019;7:205.

3. Heinig M, Adriaens ME, Schafer S, van Deutekom HWM, Lodder EM, Ware JS, et al. Natural genetic variation of the cardiac transcriptome in non-diseased donors and patients with dilated cardiomyopathy. Genome Biol. 2017;18:170.
